# Ophthalmic acid is a bloodborne metabolite that contributes to age-induced cardiomyocyte hypertrophy

**DOI:** 10.1101/2024.08.08.607218

**Authors:** Melod Mehdipour, Sangsoon Park, Wei Wei, Jonathan Z. Long, Guo N. Huang

## Abstract

Cardiac aging involves the development of left ventricular hypertrophy alongside a decline in functional capacity. Here, we use neutral blood exchange to demonstrate that the acute removal of age-accumulated blood factors significantly regresses cardiac hypertrophy in aged mice. The reversal of hypertrophy was not attributed to age-associated hemodynamic effects, implicating a role of blood-derived factors. In addition, the overarching paradigm of systemic aging maintains that the age-related overabundance of plasma proteins are largely responsible for causing pathological phenotypes in tissues. Our results suggest that blood metabolites, not proteins, drive cardiac hypertrophy instead. Upon analyzing serum metabolomics data sets, we identified ophthalmic acid as a circulating metabolite whose levels increase with advanced age. Treatment of adult mouse and neonatal rat cardiomyocytes in culture with ophthalmic acid increased their relative surface areas. This study uncovers a non-protein metabolite that may contribute to cardiomyocyte hypertrophy during aging. Identifying a method to counteract ophthalmic acid’s hypertrophic effects may reveal novel therapeutic opportunities for cardiac rejuvenation.

## Introduction

Systemic aging has been of great interest since bloodborne factors were shown to directly influence the health, regenerative capacity, and function of numerous tissues^1–5^. Old blood factors were shown to hamper tissue regeneration in young organs, whereas an old animal’s exposure to a young systemic milieu improved organ health. It was therefore proposed that young blood factors hold therapeutic and translational potential. Heterochronic blood exchange experiments have demonstrated that the influence of young blood alone can rapidly rejuvenate liver and muscle of aged mice. Hippocampal neurogenesis and cognitive performance of the aged partners, however, were not significantly affected. Young mice on the other hand exhibited a rapid and significant decline in cognition, hippocampal neurogenesis, myogenesis, and hepatogenesis^6^.

Another body of evidence suggests that systemic factors profoundly influence these mechanisms. Recently published work demonstrated that the dilution of aged mouse plasma and its replacement with a physiological solution comprised of 5% mouse serum albumin in normal saline significantly rejuvenates multiple tissues. This process termed neutral-age blood exchange was remarkable for improving liver health, muscle health, cognition, and hippocampal neurogenesis in aged mice. Neutral blood exchange concomitantly abrogated inflammation and the load of senescent cells in the aged brain^7,8^. Plasma dilution by neutral blood exchange also boosted the expression of pro-regenerative factors upon the dilution of inhibitory protein factors^7,8^.

It is thought that serum factors can influence the mammalian cardiovascular aging process. The human population is aging at a much faster rate than what was previously observed. Aging is the most significant risk factor for heart failure and the proportion of older adults who are expected to develop cardiovascular disease is increasing. Heart failure often involves cardiac hypertrophy and affects approximately 1-2% of aged adults overall, with the incidence increasing several folds in individuals who are over the age of 75. The global population of aged individuals is also expected to increase drastically over the next few decades. This emphasizes the significance of the cardiovascular system to aging^9^. Loffredo et al. demonstrated that youthful blood factors attenuated age-related cardiac hypertrophy, and that GDF11 is responsible for this effect^4^. The findings involving GDF11 were controversial as multiple groups could not replicate these results^10–12^. In addition, it is imperative to develop rapidly translatable approaches to rejuvenate the aged cardiovascular system. Considering the potential benefits of plasma dilution, we sought to determine whether a single session of neutral blood exchange can reverse age-associated cardiac hypertrophy.

Small animal blood exchange is a labor intensive and time-consuming procedure. To minimize animal distress and expenditure of resources, neutral blood exchanges were performed as blood transfusions through the retro-orbital sinuses of mice (**Figure 1**).

**Figure 1.**
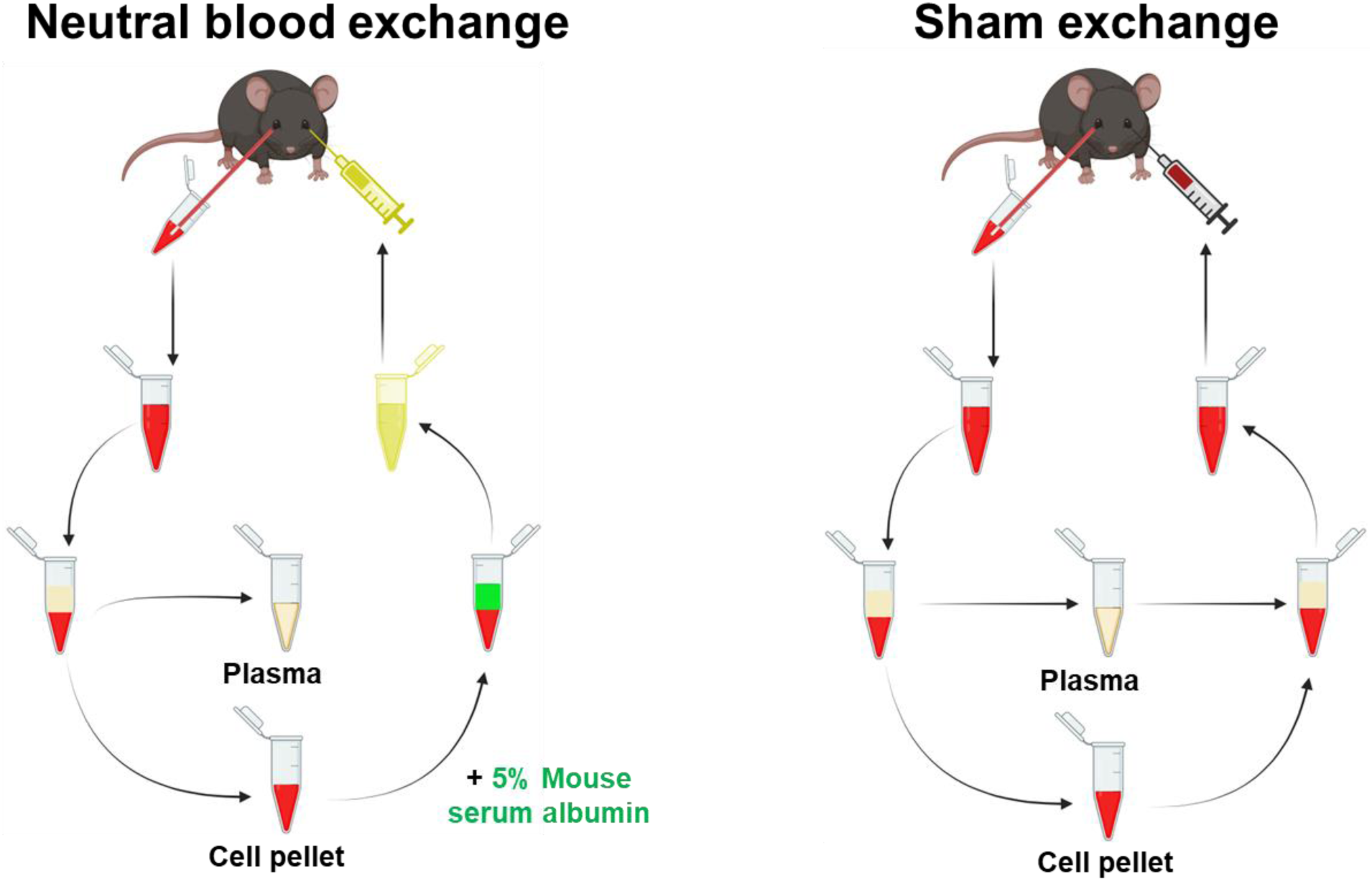
Modified Neutral-Age Blood Exchange Procedure. Young (1.5-2 months) and old (20-21 months) male C57/B6 mice underwent neutral blood exchange or sham transfusions as indicated below. Transfusions were performed such that ∼33% of platelet-rich plasma of aged mice has been replaced with 5% mouse serum albumin. Blood was drawn from the left retro-orbital sinus with a heparinized capillary tube. After gentle centrifugation (300 g for 5 min), the platelet-rich plasma fraction was decanted and replaced with an equal volume of 5% mouse serum albumin in normal saline. The blood mixture was then injected into the right retro-orbital sinus. For sham control procedures, blood from young or old mice was drawn from the left retro-orbital sinus, centrifuged, reconstituted, and then injected into the right retro-orbital sinus of the same respective mice. 6 days after exchange, mice from each of the conditions were sacrificed and their hearts were harvested for analysis. Image was created with Biorender.com.

## Results

The initial work on rejuvenating the aged heart through heterochronic parabiosis illustrated that the hearts of aged partners were reduced in size^4^. This strongly suggests that young systemic factors may play a role in reversing age-related hypertrophy. However, the effect of shared organs and environmental enrichment may play a confounding role in this phenomenon^6,7,18^. In order to address this, we sought to develop an *in vitro* cardiac hypertrophy model to determine whether factors from aged mouse serum were sufficient to cause cardiac hypertrophy and whether young serum factors can abrogate this effect.

The striking effect of neutral blood exchange on aged mouse hearts was rapidly apparent in just 6 days after the procedure (**Figure 2A**). Hearts from aged mice whose plasma was replaced with ∼33% of 5% mouse serum albumin in saline were noticeably smaller than the hearts of mice who underwent sham exchange. This observation was verified by a significant reduction of the heart weight to tibia length ratio in mice who underwent neutral blood exchange compared to the sham control group (**Figure 2B**). In old mice after neutral blood exchange, heart weight/tibia length ratios exhibited a ∼14.62% reduction in size when compared to hearts of old mice after sham operation but remained ∼24.29% larger than those of young sham-operated hearts (**Figure 2C**). We decided to normalize cardiac mass to tibia length because it is a standard method that takes body frame into account upon weighing hearts^4,19^. Though we did not observe a significant difference in body weights before and after the neutral blood exchange or sham procedures (**Supplementary Figure 1**), this method is more appropriate than normalizing to body weight since mice accumulate adipose tissue with advanced age^4,20,21^. Here, we report a ∼14.5% reduction in heart weight relative to tibia length in aged mice who have undergone neutral blood exchange.

**Figure 2.**
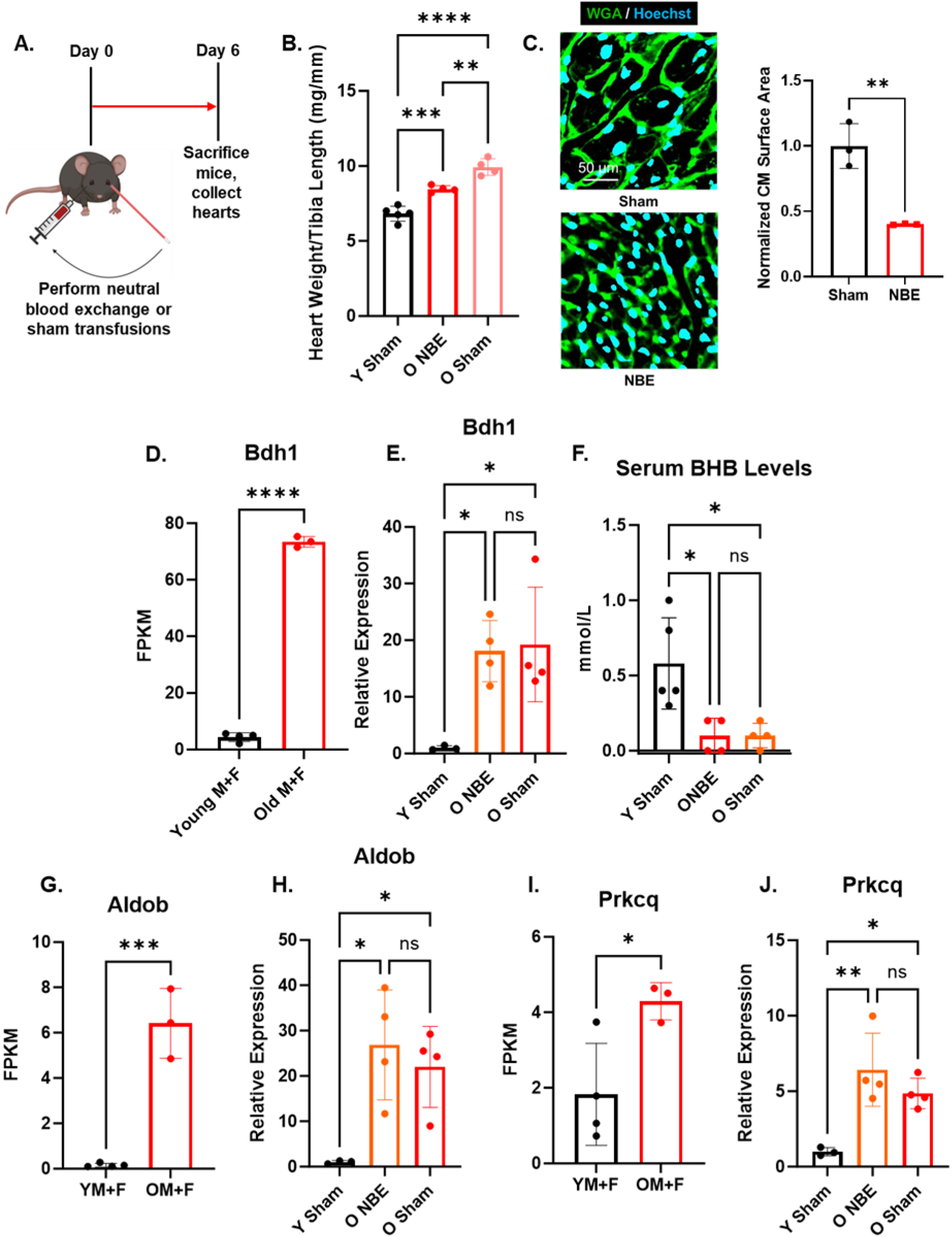
Neutral blood exchange (NBE) ameliorates age-associated cardiac hypertrophy *in vivo* independently of *Bdh1*, *Aldob*, or *Prkcq* gene activity. **A.** Schematic of the experimental procedure. Mice received NBE or sham transfusions on Day 0 and sacrificed on Day 6. **B.** Graph representing heart weight/tibia length ratios of aged mice 6 days after NBE or sham transfusions. The heart weights of the NBE group were significantly lower than those of the sham group. N = 5 for the young mouse sham exchange group (Y Sham), N = 4 for old mice who underwent neutral blood exchange (O NBE) & old mice who underwent sham exchange (O Sham), ** = P < 0.01, *** = P < 0.005, *** = P < 0.001 by One-Way ANOVA, Tukey posthoc. **C.** Wheat germ agglutinin (WGA) staining of left ventricles 6 days after NBE or sham. Representative images from each cohort as shown on the Left panel (Scale Bar = 50 μm). Quantification of the WGA staining is on the Right panel. *In situ* cardiomyocyte size was determined by measuring the cross-sectional areas of 300 – 900 myocytes per animal. Results were normalized to the sham control group and were obtained from 3 animals per group. ** = P < 0.01 by non-paired Student’s t-test. **D.** Graph representing FPKMs of Bdh1 from each age group. There is a reported ∼14-fold increase of Bdh1 with age. **** = P < 0.001 by non-paired Student’s t-test. **E.** Relative Bdh1 mRNA expression levels from the hearts of young (1.5 months), O NBE (22 months), and O sham (22 months) mice. There was an observed ∼15-17-fold increase in Bdh1 levels with age, however, NBE did not affect this age-related upregulation. * = P < 0.05 by One-Way ANOVA with Tukey posthoc. **F.** Serum BHB concentrations for each of the indicated cohorts. There was a ∼4-fold reduction in BHB with age and NBE did not appear to alter BHB concentration in aged mouse serum. * = P < 0.05 by One-Way ANOVA with Tukey posthoc. **G.** Graph representing RNAseq FPKMs of Aldob from each age group. There is a reported ∼6-fold increase of Aldob with age. *** = P < 0.005 by non-paired Student’s t-test. **H.** Relative Aldob mRNA expression levels of young (1.5 months), O NBE (22 months), and O sham (22 months) mouse hearts. Despite there being a ∼20-fold increase in Aldob with age, NBE did not affect its expression level either. * = P < 0.05 by One-Way ANOVA with Tukey posthoc. **I.** Graph representing RNAseq FPKMs of Prkcq from each age group. There is a reported ∼2-fold increase of Aldob with age. * = P < 0.05 by non-paired Student’s t-test. **J.** There was an approximate 5-fold age-associated increase in Prkcq mRNA expression levels in the hearts of young (1.5 months), O NBE (22 months), and O sham (22 months) mice. Panel **A** was created with Biorender.com.

We then determined whether the apparent reduction of cardiac hypertrophy resulted from changes in cellular size within the heart. We performed histological analysis by staining paraffinized heart sections with wheat germ agglutinin conjugated to a green fluorophore. We found a significant reduction in cardiomyocyte surface area in hearts from neutral blood exchange mice compared to the sham group (**Figure 2B**). Upon quantification, the relative cardiomyocyte size reduction was approximately two-fold. The above results appear to be consistent with the effect of heterochronic parabiosis on the aged heart^4^. Thus, the dilution of aged blood plasma appears to be sufficient for reversing the hypertrophic cellular phenotype in old mouse hearts.

In effort to elucidate a molecular mechanism, we consulted a bulk RNAseq dataset that compared differential gene expression changes in the hearts of mice spanning from 1 month of age to 22 months of age^22^. We compared the fragments per kilobase of transcript per million fragments mapped (FPKM) of 22-month-old mice in relation to the logarithm of the FPKM fold change of 22-month-old mice to 1 month old mice (**Supplementary Figure 2A**). We then utilized the following criteria to select potential candidates for further investigation: the FPKM of the upregulated gene must be at least 10, the fold-change of genes compared between 22 month and 1 month old mice must be at least 2, and the relationship between the compared genes must be statistically significant (P-value less than 0.05). We selected genes that were upregulated in order to focus on aging-induced factors that may give rise to age-dependent cardiac hypertrophy. β-hydroxybutyrate dehydrogenase 1 (Bdh1) was found to fit the criteria and was subsequently validated by RT-qPCR (**Figure 2D-E**). Since RNAseq data set investigated the differences between 1-month-old vs 22-month-old mice, we performed RT-qPCR to verify Bdh1 expression levels in young adult mice of 2 months of age vs 21-month-old animals. We observed an age-related increase in this instance; however, the fold change is not as pronounced (**Supplementary Figure 2B).** These results suggest that Bdh1 expression levels may be gradually increasing throughout the mouse’s lifetime.

Bdh1 was not previously shown to increase in the context of age-related cardiac hypertrophy. However, studies suggest that Bdh1 is upregulated at the mRNA and protein levels during transverse aortic constriction-induced pathological cardiac hypertrophy^23,24^. It is also known that the capacity of fatty acid oxidation is reduced in the aged heart, and it is speculated that increased ketone utilization may compensate for this^25^. Ketone bodies are used as an efficient energy source in failing hearts and its supplementation is hypothesized to be beneficial^26^. It is also thought that beta-hydroxybutyrate (BHB), a substrate of Bdh1, is used by failing hearts as a stress defense^24^. We therefore reasoned that increased levels of Bdh1 may be associated with age-related cardiac hypertrophy.

We sought to investigate whether the apparent reduction of heart weight in mice who have undergone neutral blood exchange exhibited a concomitant reduction of *Bdh1* mRNA expression. To our surprise, *Bdh1* levels in hearts from old neutral blood exchange mice remained nearly as elevated as that of old sham mice (**Figure 2E**). Despite the lack of reduction in *Bdh1* expression levels, we measured serum BHB levels to determine whether neutral blood exchange may still increase its concentration. There was a sharp age-related decline in serum BHB concentration, but neutral blood exchange appeared to have no effect on this (**Figure 2F**). Neutral blood exchange may reduce age-related cardiac hypertrophy by a mechanism that is independent of Bdh1-BHB metabolism. Aldolase B and Protein Kinase C Theta were found to have also exhibited a robust age-related increase in mRNA expression, but their up-regulation was unaffected by neutral blood exchange (**Figure 2G-J**).

One concern regarding any potential causes for age-associated cardiac hypertrophy are hemodynamic effects related to blood pressure. To investigate the hemodynamic phenomenon between young (2-3 months) and aged (22-23 months) mice, we measured their blood pressure with a computerized tail cuff system (BP-2000, Visitech Systems, Apex, NC). We saw no significant differences in systolic or diastolic pressure in young or aged mice (**Supplementary Figure 3A-B**). There were also no differences in heart rate either (**Supplementary Figure 3C**). These data suggest that blood pressure or heart rate do not play a significant role in the induction of age-associated cardiac hypertrophy.

We then sought to detect potential changes in cardiac hypertrophy markers brain natriuretic peptide (BNP), atrial natriuretic peptide (ANP), and sarcoplasmic reticulum calcium ATPase (SERCA-2) using the same primers as those from a previously published study^4^. We did not detect any significant age-related changes in transcript levels for either of the genes. Neutral blood exchange transfusion did not significantly alter transcript levels either (**Supplementary Figure 4A-C**). An independent RNAseq dataset has also confirmed the apparent lack of age-related changes in these gene targets^22^ (**Supplementary Figure 2**). These results underscore the need to identify robust cardiac markers that change with age.

The results from **Figure 2** suggest that serum factors may play a direct role in modulating age-related cardiac hypertrophy. We therefore sought to test potential hypertrophic effects of aged mouse serum on adult mouse cardiomyocytes. We devised an *in vitro* paradigm where adult mouse cardiomyocytes were cultured with standard culture medium, 50% young mouse serum, 50% aged mouse serum, or 10 μM Isoproterenol. Isoproterenol is β-adrenergic receptor agonist that is known to increase heart rate and contractility through a G-protein coupled receptor-dependent mechanism^27^. It is also known to induce cardiac hypertrophy. Cardiomyocytes treated with isoproterenol exhibit an average surface area increase by ∼10-20%^28,29^. After incubating adult mouse cardiomyocytes with serum, isoproterenol, or negative medium for 24 hours, cardiomyocytes were fixed, and the average relative surface area of individual cardiomyocytes was obtained (**Figure 3A**).

**Figure 3.**
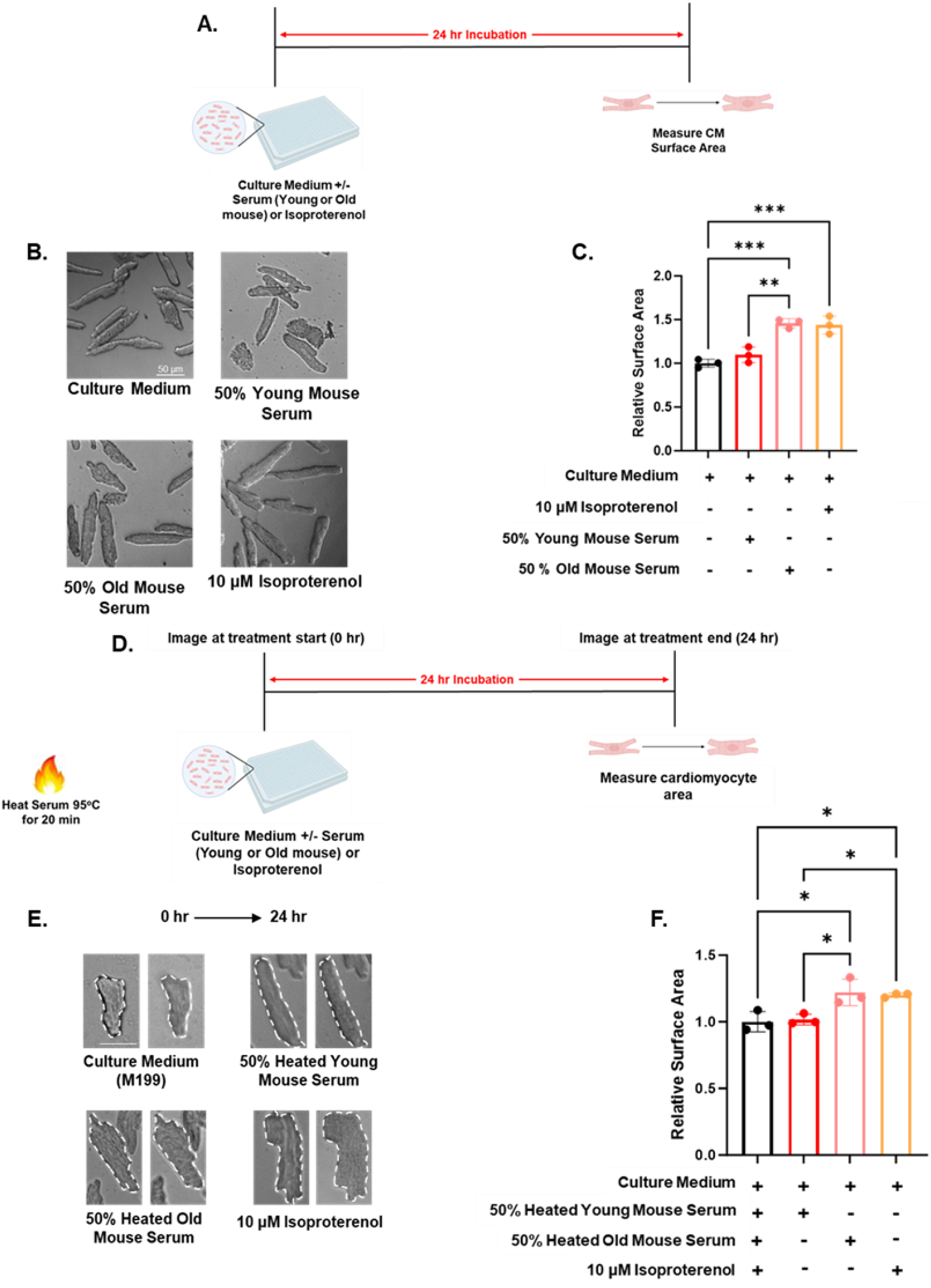
Blood heterochronicity of non-protein factors influences cardiomyocyte size *in vitro*. **A.** Adult mouse cardiomyocytes were freshly isolated and seeded on a 384 well plate. They were incubated in standard culture medium, treated with 50% young (2 months) mouse serum, 50% old (22 months) serum, and 10 μM isoproterenol. Cardiomyocytes were incubated in their respective treatment media for 24 hours and imaged on brightfield. **B.** Representative images of cardiomyocytes from each cohort. **C.** Cardiomyocytes treated with old mouse serum exhibited a significant increase in relative surface area compared to those treated with young mouse serum. Average surface areas were obtained from ∼200-300 cardiomyocytes per cohort. N = 3 in technical replicates per cohort. ** = P < 0.01 & *** = P < 0.005 by One-Way ANOVA, Tukey posthoc. Scale bar = 50 μm. Data shown as mean +/-Stdev. **D.** Experimental schematic. Sera from young (2-3 months) and old (20-22 months) mice were heated for 20 minutes at 95°C, centrifuged at 21,130 g, and the remaining supernatant was used to treat adult mouse cardiomyocytes at a concentration of 50% (v/v). **E.** Representative images taken immediately at the beginning of treatment (0 hr) and after 24 hr of incubation (24 hr). **F.** Relative to culture medium condition, the heated fraction of aged serum induced hypertrophy to a similar extent of the positive control, 10 μM Isoproterenol. *p < 0.05 by One-Way ANOVA with a Tukey post-test, all experiments were performed with n = 3 per cohort in technical triplicate.

The relative surface area of adult mouse cardiomyocytes treated with isoproterenol was significantly increased as expected. Adult mouse cardiomyocytes treated with 50% young mouse serum did not exhibit a significant surface area size when compared to adult mouse cardiomyocytes incubated with standard culture medium. Cardiomyocytes that were treated with 50% aged mouse serum exhibited a significant increase in surface area size that was comparable to that of adult mouse cardiomyocytes treated with isoproterenol (**Figure 3B-C**). 50% young and 50% aged sera increased the size of P2 rat cardiomyocytes to a similar degree that was not statistically significant (**Supplementary Figure 5A-B**). This phenotype persists even when neonatal rat cardiomyocytes are treated with young or old serum concentrations as low as 3% (**Supplementary Figure 5C**).

Blood serum contains a plethora of bioactive protein and non-protein molecules. We aimed to elucidate this by examining the effects of serum or heated serum from young or old mice. We devised an *in vitro* adult mouse cardiomyocyte hypertrophy platform to accomplish this. Freshly isolated adult mouse cardiomyocytes were treated with young mouse serum, aged mouse serum, isoproterenol, or negative culture medium for 24 hours. Cardiomyocytes were then fixed and imaged for cell surface area analysis (**Figure 3D**). To determine whether proteins or non-protein species may be responsible for this effect, we boiled young and old mouse sera at 95°C for 20 minutes. Potential bioactivity of the remaining supernatants was assayed on adult mouse cardiomyocytes. We confirmed size increases of representative cardiomyocytes by comparing individual cells immediately at the beginning of treatment (0 hr) and 24 hours after treatment (**Figure 3E**). Interestingly, aged mouse sera retained its hypertrophic effect after boiling (**Figure 3F**).

The above results strongly suggest that a heat-resistant small molecule metabolite may be responsible for eliciting a hypertrophic effect. A recent metabolomics dataset reported age-related changes of circulating metabolites in the sera of mice^30^. A tripeptide named ophthalmic acid was identified to be among the most upregulated metabolites in the aged serum (**Figure 4A**). Ophthalmic acid is a tripeptide molecule that bears structural similarity to glutathione. The peptide sequences of ophthalmic acid and glutathione are nearly identical, however, the second amino acid in ophthalmic acid is an aminobutyrate residue in place of cysteine that is found in glutathione. Interestingly, both ophthalmic acid and glutathione are synthesized by the same enzymes. It was initially thought that ophthalmic acid is an accidental byproduct of glutathione synthesis that lacks a coherent biological function. However, recent studies suggest that ophthalmic acid acts to negatively regulate glutathione activity. The direct effects of ophthalmic acid on tissue function remain largely unknown^31,32^.

**Figure 4.**
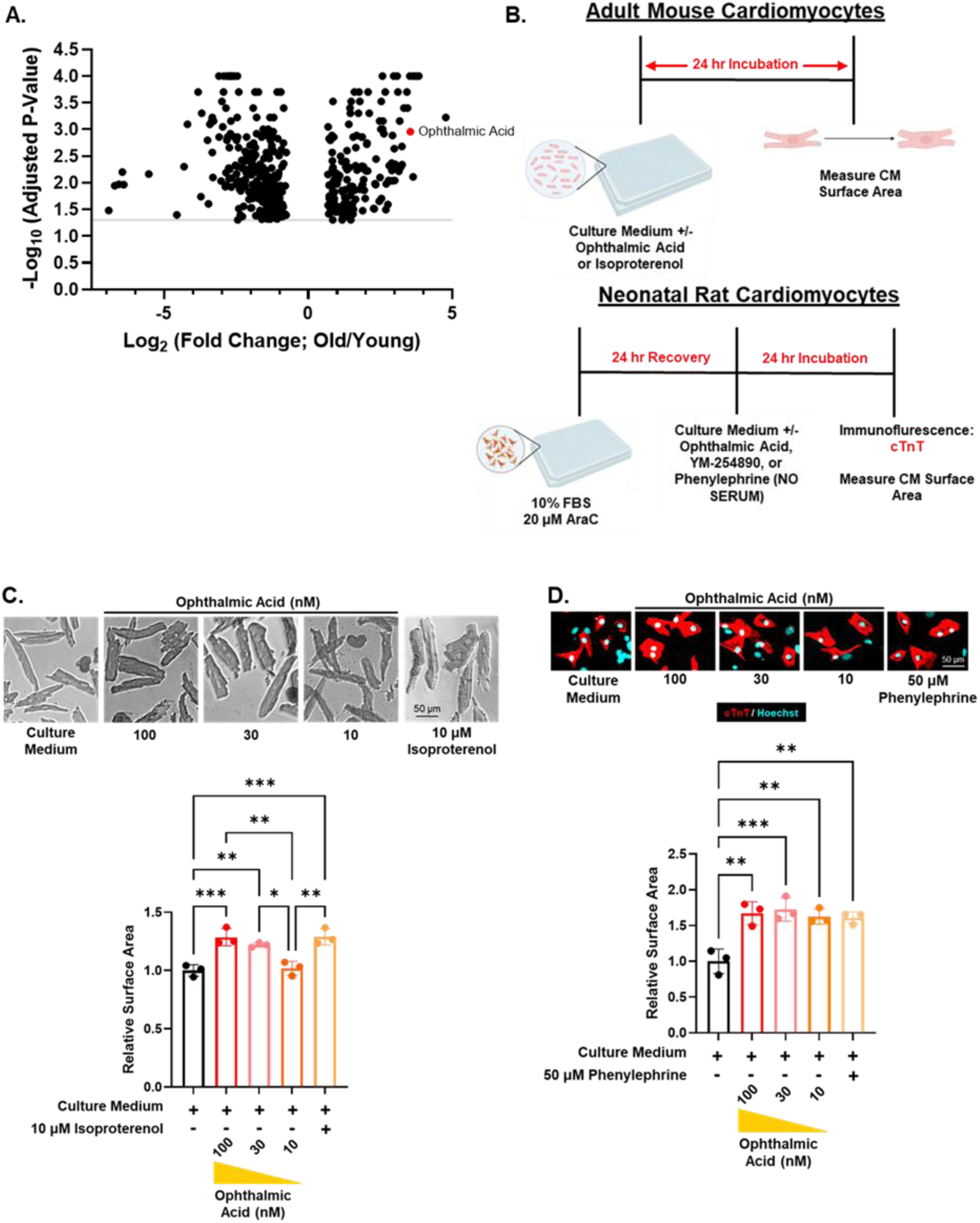
Ophthalmic acid causes cardiac hypertrophy in vitro. **A.** Volcano plot of upregulated and downregulated metabolites in aged mouse blood sera relative to young levels. The gray line indicates an adjusted P-value that is equal to 0.05. **B.** Treatment paradigms for adult mouse cardiomyocytes and neonatal rat cardiomyocytes. **C.** 30 nM of ophthalmic acid appears to be sufficient to induce hypertrophy in adult mouse cardiomyocytes. **D.** Ophthalmic acid induces hypertrophy in neonatal rat cardiomyocytes but at lower doses. cTnT = cardiac troponin T. *p < 0.05, **p < 0.01, and **p < 0.005 by One-Way ANOVA with a Tukey post-test, all experiments were performed with n = 3 per cohort in technical triplicate.

To determine whether this metabolite can induce cardiomyocyte hypertrophy directly, we treated adult mouse cardiomyocytes and neonatal rat cardiomyocytes with varying concentrations of ophthalmic acid. We initiated the experiment with a concentration of 30 μM, progressively titrating the dosage downwards by a factor of 3. Cardiomyocytes were treated with culture medium only, ophthalmic acid, or a hypertrophic stimulant (isoproterenol or α-adrenergic agonist phenylephrine) for 24 hours in serum-free conditions. Of note, adult mouse cardiomyocytes were treated with compounds on the same day as isolation. Neonatal rat cardiomyocytes were allowed to recover for 24 hours after isolation, and then treated with compounds for an additional 24 hours (**Figure 4B**). It was observed that 30 nM of ophthalmic acid was sufficient to induce hypertrophy in adult mouse cardiomyocytes after following this paradigm (**Figure 4C**). Interestingly, concentrations as few as 10 nM still induced hypertrophy in neonatal rat cardiomyocytes (**Figure 4D**). The variance in sensitivity to ophthalmic acid could be attributed to the higher inclination of neonatal rat cardiomyocytes towards growth.

## Discussion

Aging is the largest risk factor for developing cardiovascular diseases because the prevalence of such pathologies drastically increases as one grows older^33^. Due to an apparent increase in average human lifespan worldwide, it has been predicted that at least one in four individuals will be 65 years or older in the next 10 – 15 years^34,35^. Left ventricular hypertrophy is a significant feature of cardiac aging which contributes to diastolic dysfunction and heart failure^34,36,37^. Cardiac hypertrophy is also thought to be a compensatory response that results from cardiomyocyte loss with advancing age^34,38^. Despite the presence of conflicting reports, current evidence suggests that left ventricular thickening affects both sexes instead of mostly males^34,36,39,40^. Cardiac hypertrophy may also contribute to other detrimental processes such as arrhythmias, fibrosis, electrophysiological derangements, or heart failure^34,41^. These findings imply that cardiovascular aging may be regulated by distinct pathological mechanisms.

The central hypothesis of our study is that the hypertrophic phenotype of the aged heart is reversible after acute plasma dilution. This hypothesis was tested by replacing ∼33% of aged mouse plasma with an “ageless” physiologic solution as described previously^7,8,15,42^. C56BL/6 mice were selected for these experiments since they exhibit cardiac aging phenotypes that are similar to those of humans^4^. We have indeed found that systemic factors may contribute to increased average cardiomyocyte size. We also observed that cardiac mass was rapidly reduced after a single neutral blood exchange procedure. These results compound the recent paradigm which maintains that the ectopic administration of young blood factors may not be necessary to perpetuate rejuvenating phenotypes.

It was proposed that the levels of Growth Differentiation Factor 11 (GDF11) decrease in the blood of aged mice and that exogenous supplementation in old animals reduced cardiac hypertrophy^4^. Though encouraging, these findings are fraught with controversy. One study has demonstrated that GDF11 can exhibit anti-hypertrophic effects in cardiac tissue through SMAD2/3 signaling^43–45^. It was also reported that targeted GDF11 delivery to the heart can increase the number of cardiac stem cells in older animals with an associated improvement in cardiac function^43,46^. Other reports, however, were unable to replicate the beneficial findings. One group used a similar dosing scheme to that of Loffredo et al^4^, but they found that this dose was not sufficient to alter cardiomyocyte surface area *in vivo*^12^. This same group^47^ eventually found that GDF11 doses that were 10-fold or 50-fold higher than what was initially used^4^ conferred anti-hypertrophic effects to the heart. However, these doses were shown to be directly associated with cachexia and death^47^.

The results of our present study suggest that factors other than “youthful” proteins may be responsible for the apparent reduction of cardiac hypertrophy. It is presumed that the dilution effect model could in part contribute to this phenotype^7,8^, but further study is needed to confirm this notion.

We also show that hemodynamics is unlikely to explain any changes to heart weight with increased age. Apparent changes in the mRNA expression of ANP, BNP, and SERCA-2 may also be unlikely to account for the reduction of hypertrophy by neutral blood exchange. Though these markers may be indicative of cardiac hypertrophy to an extent, we did not observe differential expression of these genes in relation to age, and this was apparent in the study reported by Loffredo et al^4^. We also noticed substantial variability in the expression of these markers across each sample per cohort. Lastly, a bulk RNAseq dataset strongly associates the age-related upregulation of Bdh1 in heart tissue^22^.

We were able to replicate these findings by RT-qPCR, but its increased expression in aged mice was unabated by neutral blood exchange. Serum β-hydroxybutyrate levels were found to decrease with age, suggesting that increased Bdh1 expression is potentially compensating for a metabolic perturbation in the aged heart. NBE did not increase serum BHB in aged mice alluding to the possibility that ketone body metabolism may not play a direct role in the reduction of hypertrophy. Functional studies and the elucidation of a molecular mechanism will require further investigation.

## Acknowledgements

We thank the members of the Huang lab for their discussions. This work is supported by a T32 training grant (to M.M.), NIH Awards (R01HL138456 and R01HL157280), Department of Defense (W81XWH1910206), Tobacco-Related Disease Research Program Award P0558275, American Heart Association Established Investigator Award, American Federation for Aging Research Junior Faculty Grant, BAKAR Aging Research Institute Investigator Award (G.N.H.).

## Materials and Methods

### Adult Mouse Cardiomyocyte Isolation, Plating, & Hypertrophy Assay

Adult mouse cardiomyocytes were isolated as previously described (Judd 2015). Wells of a 384 well plate (CELLTREAT Scientific Products, Pepperell, MA, USA, Lot #210612-348) were pre-coated with 5 μg/mL of α-laminin for at least one hour at 37°C. Freshly isolated adult mouse cardiomyocytes suspended in plating medium were seeded in pre-coated wells at a density of ∼3000 cells/cm^2^ and 50 μL per well. Cardiomyocytes were incubated for one hour in plating medium. The plating medium was then exchanged with culture medium and/or treatment medium as indicated. The negative medium condition was comprised of standard adult mouse cardiomyocyte culture media. Young and old mouse sera conditions contained a 1:1 formulation of 2X culture medium and serum. Isoproterenol (Isoprenaline hydrochloride, Sigma-Aldrich, St. Louis, MO, USA, Lot #BCBZ5101) was dissolved to a final concentration of 10 μM in culture medium. A stock concentration of ophthalmic acid (Millipore Sigma, Cat#91351-10MG) was dissolved to 10 mg/mL (approximately 34.6 mM) and diluted to the indicated working concentrations.

Cardiomyocytes were incubated in treatment media for 24 hours. Cardiomyocytes were imaged using a Nikon Eclipse Ti microscope with Nikon DS-Qi2 camera at 10X magnification. Brightfield images of the cardiomyocytes were then acquired at 10X magnification and the surface area of each cardiomyocyte per condition was recorded. Average surface areas were obtained from ∼200-300 cardiomyocytes per cohort and were normalized to those of the negative medium condition.

### Neonatal Rat Ventricular Cardiomyocyte Isolation, Plating, and Hypertrophy Assay

Formulations of relevant buffers:

- *Cardiomyocyte Isolation Buffer:*

- *Perfusion Buffer:* Cardiomyocyte isolation buffer + 0.4 mM ethylene glycol-bis(β-aminoethyl ether)-N,N,N′,N′-tetraacetic acid - *Digestion Buffer:*

***- Cardiomyocyte Isolation Buffer:***
| **Component** | **g/500mL** |
| --- | --- |
| NaCl | 3.518 |
| KCl | 0.5479 |
| KH<sub>2</sub>PO<sub>4</sub> | 0.0408 |
| Na<sub>2</sub>HPO<sub>4</sub>·7H<sub>2</sub>O | 0.0804 |
| MgSO<sub>4</sub>·7H<sub>2</sub>O | 0.1478 |
| NaHCO<sub>3</sub> | 0.1932 |
| Taurine | 1.8765 |
| 2,3-Butanedione Monoxime | 0.5055 |
| Glucose | 0.5449 |
| HEPES | 1.1915 |

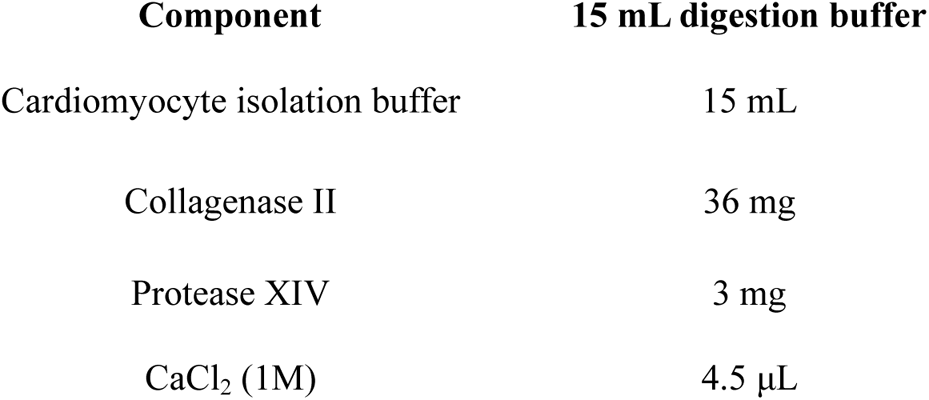
- *Stop Buffer:*

- *Wash Buffer 1:*

- *Stop Buffer:*
| Component | Volume |
| --- | --- |
| FBS | 500 μL |
| CaCl<sub>2</sub> (1M) | 1.6 μL |

- *Wash Buffer 2:*

- *Wash Buffer 1:*
| Component | Volume |
| --- | --- |
| Cardiomyocyte Isolation Buffer | 3 ml |
| FBS | 300 μL |
| CaCl<sub>2</sub> (1M) | 2.1 μL |

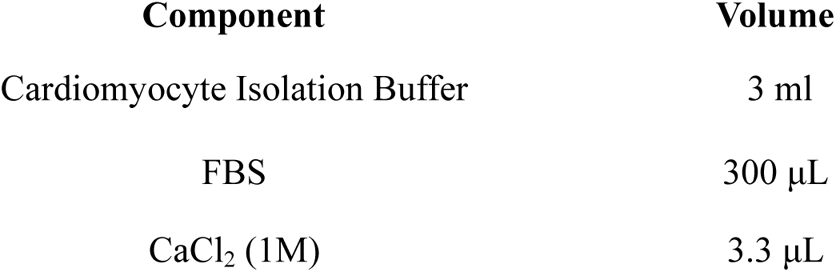 - *Culture Medium:*

### 10% fetal bovine serum + 100 U/mL penicillin-streptomycin in Dulbecco’s Modified Eagle Medium

#### Isolation

Wells of a 384 well plate (CELLTREAT Scientific Products, Pepperell, MA, USA, Lot #210612-348) were pre-coated with 1% gelatin in sterile 1x PBS for at least one hour at 37°C. 2 day postnatal Sprague Dawley rat pups were anesthetized on ice for 5 minutes and euthanized by cervical dislocation. Their hearts were dissected and the atria were removed with micro dissecting scissors. The aorta was cannulated with a blunted 25G needle and perfused with perfusion buffer and digestion buffer. Perfused hearts were then incubated in digestion buffer for 30 min - 1 hour at 37°C with periodic agitation. Stop buffer was added to the digestion mixture and undigested debris was discarded. Isolated cells were gently centrifuged at 200 g for 3 min and the supernatant was discarded. The cells were rinsed with wash buffer 1, centrifuged at 200 g for 3 min, washed once more with wash buffer 2, centrifuged at 200 g for 3 min, and resuspended in culture medium.

#### Pre-plating and plating

To remove fibroblasts, the cell suspension was pre-plated in a 6-well cell culture plate (Corning Life Sciences, Corning, NY, Cat #0720091) for 1-2 hours at 37°C. The number of live cells that were confirmed with Trypan blue (Fisher Scientific, Cat #BW17942E) stainings were quantified with a hemocytometer. 5,500 neonatal rat cardiomyocytes were seeded per well and were allowed to recover for 24 hours in the presence of 20 μM cytosine arabinoside (Sigma, Cat # C1768-1G).

#### Hypertrophy assay and immunostaining

Hypertrophy assays were performed with young mouse serum, old mouse serum, ophthalmic acid, or phenylephrine (Sigma, Cat #P6126-10G) as indicated. Cardiomyocytes were incubated with compounds for 24 hours and fixed with 3.7% formaldehyde in 1X PBS for 10 minutes at room temperature. The cells were permeabilized and blocked with 5% normal donkey serum in 1X PBS with 0.1% Triton X-100 for 30 minutes at room temperature. Cardiomyocytes were incubated with anti-cardiac troponin T antibodies (1:500 in 1X PBS with 0.1% Triton X-100, Thermo Scientific, Lot# MS-295-P1) for 1-2 hours at room temperature. The cells were rinsed with 1X PBS with 0.1% Triton X-100 three times and incubated with AlexaFluor 555 donkey anti-mouse (1:500, Life Technologies, Cat #A31570) antibodies and Hoechst 33342 dye (10 μg/mL; R&D Systems, Cat #5117) for 30 min-1 hour a room temperature. The cells were rinsed with 1X PBS and imaged with a Nikon Eclipse Ti microscope, using a Nikon DS-Qi2 camera at 10X magnification.

### Neutral Blood Exchange or Sham Blood Exchange Procedures

All animal procedures were performed as per the strict guidelines of the Institutional Animal Care & Use Program of the University of California, San Francisco. Young (1.5-2 months) and old (20-21 months) male C57/BL6 mice were anesthetized with 3% – 5% isoflurane in supplemental oxygen.

A heparinized micro-hematocrit capillary tube (Fisherbrand, Pittsburgh, PA, USA, Cat #22-362-566) was used to apply gentle pressure to the right retro-orbital sinus and to collect approximately one-third of the mouse’s total blood volume. Blood was collected into an Eppendorf tube that contained ∼1 unit of heparin in saline. The collected blood was centrifuged at 300 g for 5 minutes. For the NBE condition, the platelet-rich plasma was decanted and replaced with an equal volume of 5% mouse serum albumin (Molecular Innovations, MI, USA, IMSALB1000MG) in normal saline.

This blood mixture was reconstituted, drawn into a syringe, and injected back into the left retro-orbital sinus of the same mouse that the blood was initially drawn from. These steps are identical for the sham condition, except that the centrifuged blood was reconstituted in its own plasma in place of the 5% mouse serum albumin solution. The mice were sacrificed 6 days after the exchanges, and their hearts were harvested for analysis.

### Wheat Germ Agglutinin Staining & Quantification

Heart samples were incubated in 4% paraformaldehyde for 48 hours at 4°C. The hearts were then submerged in 70% ethanol and shipped to AML Laboratories (St. Augustine, FL, USA) for paraffin embedding.

Paraffinized hearts were sectioned with a microtome at a thickness of 5 μm. The heart sections were gradually rehydrated by a successive wash series using the following solutions in the order presented: xylenes (2 min), equal parts of xylenes and 100% ethanol (2 min), 95% ethanol (2 min), 70% ethanol (2 min), 50% ethanol (2 min), Milli-Q ultrapure water (2 min). The slides were then incubated with Wheat Germ Agglutinin Oregon Green Conjugate 488 (Life Technologies, Carlsbad, CA, USA, Lot #1566514) at a 1:200 dilution in phosphate buffered saline for 30 minutes at room temperature (∼25°C). The slides were rinsed 3 times with 0.2% Triton X-100 in phosphate buffered saline with 5 minutes per rinse.

Cardiomyocytes in the left ventricle were then imaged with a Nikon Eclipse Ti fluorescent microscope with a Nikon DS-Qi2 camera. Images were taken at 10x magnification and processed with Fiji. The surface areas of the cardiomyocytes were obtained and subsequently normalized to the old sham group.

### Non-Invasive Hemodynamics

We used a non-invasive tail-cuff device (BP-2000, Visitech Systems, NC, USA) to obtain systolic blood pressure, diastolic blood pressure, and heart rate simultaneously. Individual mice were trained for 3-5 consecutive days in the pre-warmed tail cuff system to acclimate them to the procedure. Measurements of blood pressure and heart rate were then obtained over the span of three additional consecutive days.

### Serum β – Hydroxybutyrate Concentration Measurements

β – hydroxybutyrate concentrations in serum were obtained with a digital Precision Xtra Blood Glucose & Ketone Monitoring System (Abbot, CA, USA, XEGZ162-P08F7) with its associated Precision Xtra blood ketone test strips (Abbot, CA, USA, X001M90E19) as per the manufacturer’s instructions. This blood ketone detection system exclusively detects β – hydroxybutyrate.

### Gene Target Validation

1 mL of TRIzol Reagent (ThermoFisher Scientific, Weltham, MA, Cat. #15596026) was added to pieces of heart apexes in Eppendorf tubes on ice. Tissues were homogenized with Qiagen stainless steel beads (Qiagen, Hilden, Germany, Lot #2271122) using the Qiagen TissueLyser II system (Qiagen, Hilden, Germany) at a frequency of 30 s^-1^ for 5 minutes at 4°C. Once 200 μL of chloroform was added to each sample, tubes were shaken vigorously for 15 seconds and allowed to sit at room temperature for 3 minutes. The samples were then centrifuged at 21,130 g for 15 minutes at 4°C. ∼400 μL of the aqueous layer was transferred to new Eppendorf tubes where 500 μL of isopropanol and 1 μL of 10 mg/mL glycogen (ThermoFisher Scientific, Weltham, MA, Cat. #2636533) were added. The samples were mixed and centrifuged at 21,130 g for 15 minutes at 4°C. The supernatant was decanted, and 1 mL of 70% ethanol was used to wash each pellet. Samples were centrifuged at 8,000 g for 5 minutes at 4°C. The supernatant was decanted and the pellets were allowed to airdry completely. RNA was quantified using the NanoDrop 2000c system (ThermoFisher Scientific, Weltham, MA).

Purified RNA samples were reverse transcribed using iScript Reverse Transcription Supermix (Bio-Rad, CA, USA, Cat #1708841). 250 ng of RNA from each sample was reverse transcribed in a final reaction volume of 5 μL. The reaction protocol was performed according to the manufacturer’s instructions. Quantitative real-time polymerase chain reaction was performed with the QuantStudio 5 Real-Time PCR System (ThermoFisher Scientific, Weltham, MA, Cat. #A28574).

The following primers were used:

*Bdh1* forward: 5’ – ACAAGACACACGCTGTTGTTT – 3’

*Bdh1* reverse: 5’ – CTCTTCAAGCTGTCCAGTTCC – 3’

*Aldob* forward: 5’ – GAAACCGCCTGCAAAGGATAA – 3’

*Aldob* reverse: 5’ – GAGGGTCTCGTGGAAAAGGAT – 3’

*Prkcq* forward: 5’ – TATCCAACTTTGACTGTGGGACC – 3’

*Prkcq* reverse: 5’ – CCCTTCCCTTGTTAATGTGGG – 3’

*Aldob* forward: 5’ – GAAACCGCCTGCAAAGGATAA – 3’

*Aldob* reverse: 5’ – GAGGGTCTCGTGGAAAAGGAT – 3’

## Figures

**Supplementary Figure 1.**
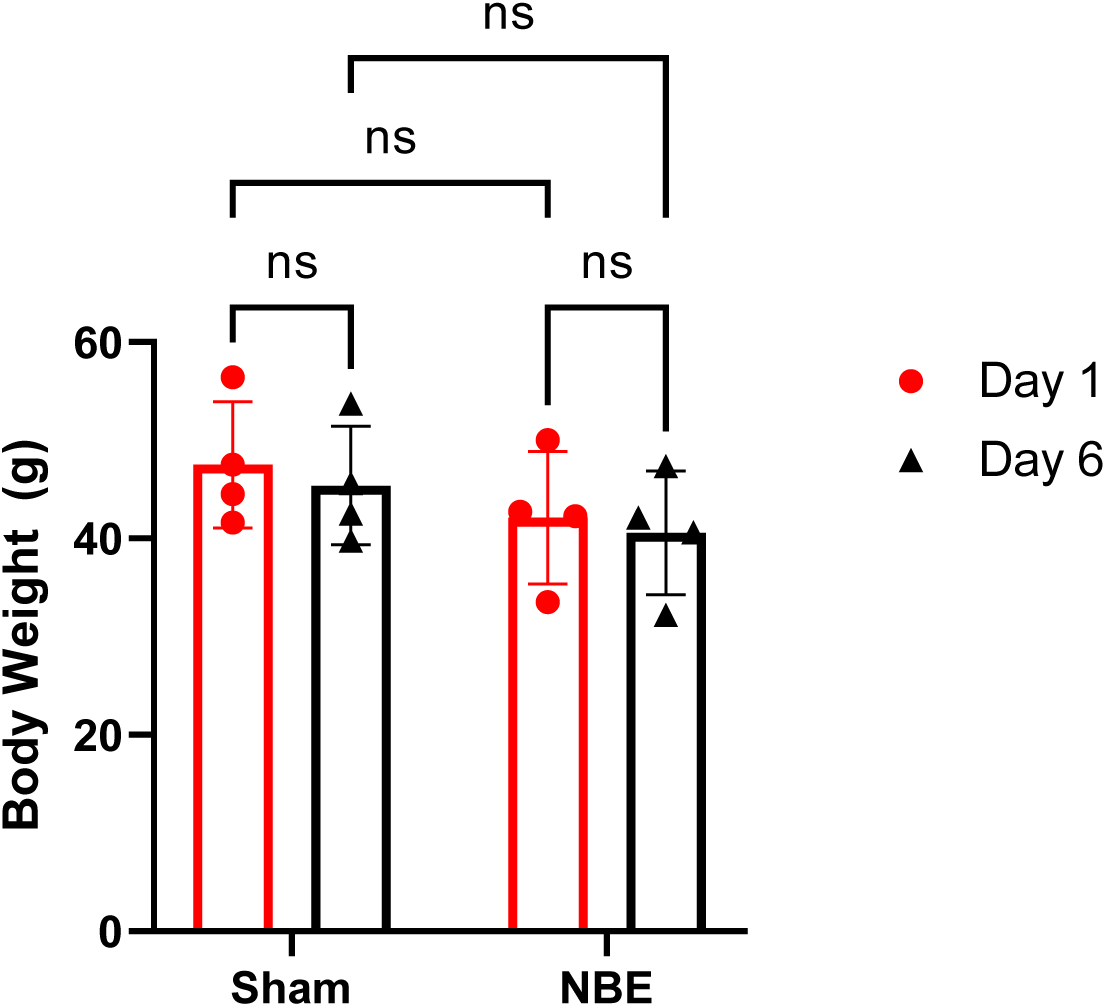
The NBE procedure did not significantly alter the body weight of mice. Aged mice underwent NBE or sham transfusion on Day 0. Their body weights were obtained the following day on Day 1, and just prior to sacrifice on Day 6.

**Supplementary Figure 2.**
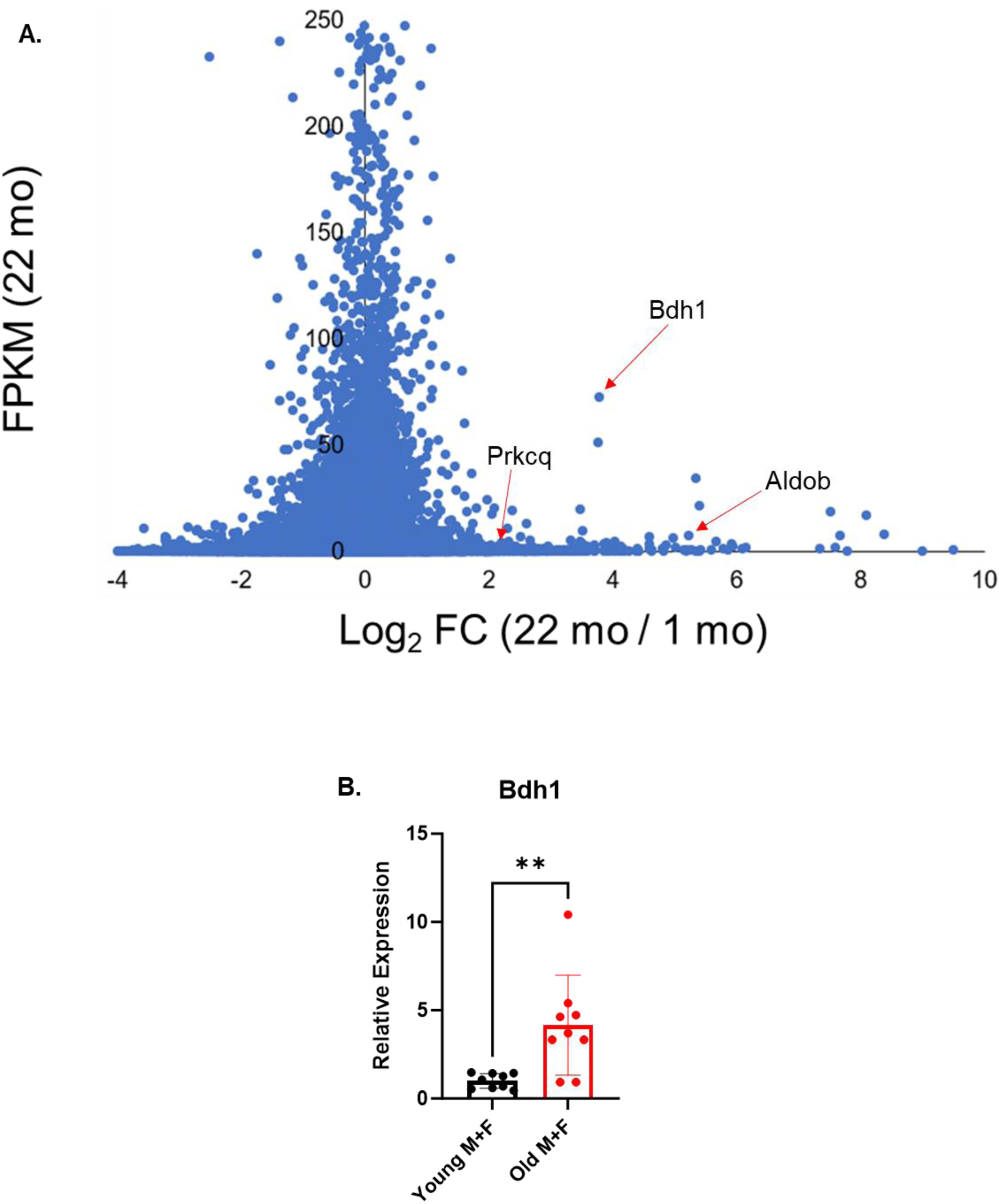
**A.** FPKM vs Log_2_ fold change (FC) plot of the bulk RNAseq data. The positions of *bdh1*, *aldob*, and *prkcq* is indicated with the red arrow. Each data point on the plot represents the means of three 22-month-old mice and four 1-month-old mice. **B.** RT-qPCR of Bdh1 levels for 2-month-old vs 21-month-old male and female (M+F) mice. ** P < 0.01 by Student’s t-test, n = 9 biological replicates per cohort.

**Supplementary Figure 3.**
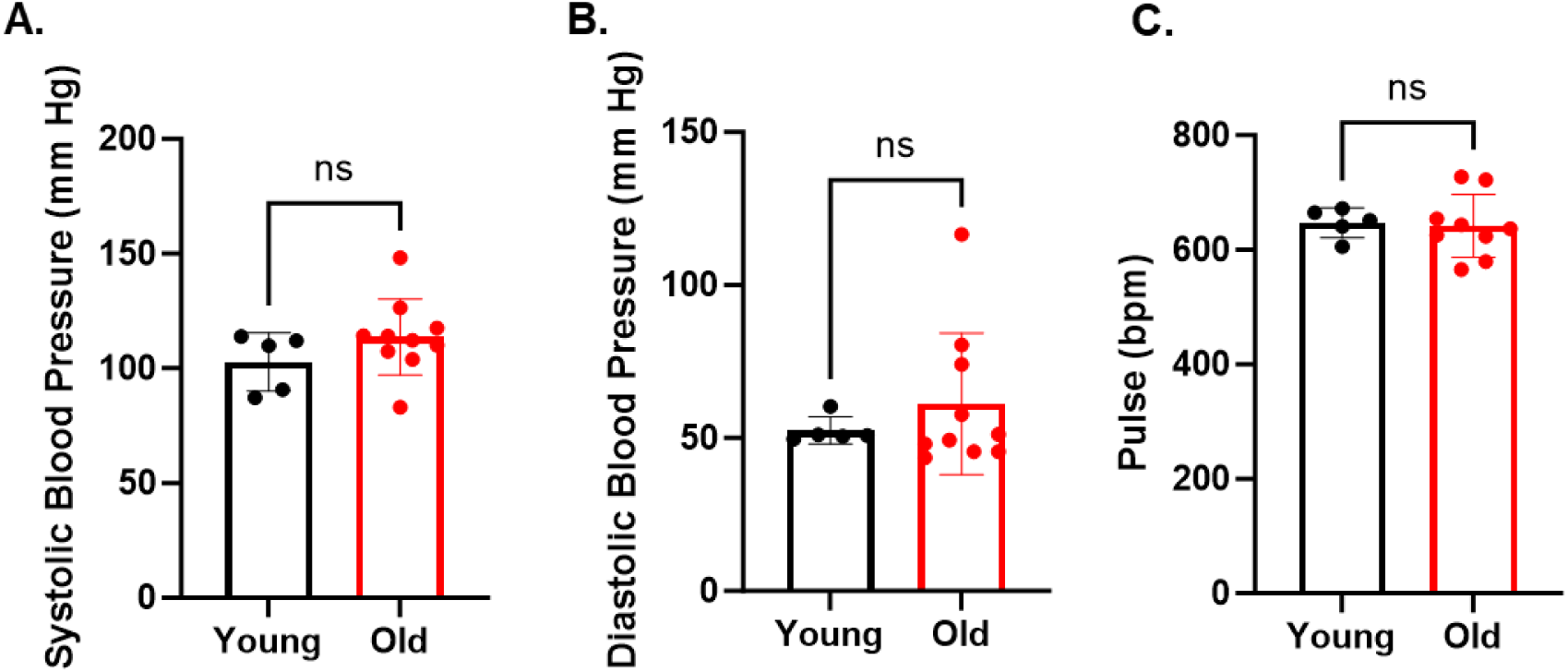
Age-related hypertrophy cannot be explained by hemodynamics. The computerized tail cuff system was used to obtain (**A**) systolic blood pressure, (**B**) diastolic blood pressure, and (**C**) pulse. There were no statistically significant differences between young (n = 5) and old (n = 10) mice. P = ns by unpaired Student’s t-test.

**Supplementary Figure 4.**
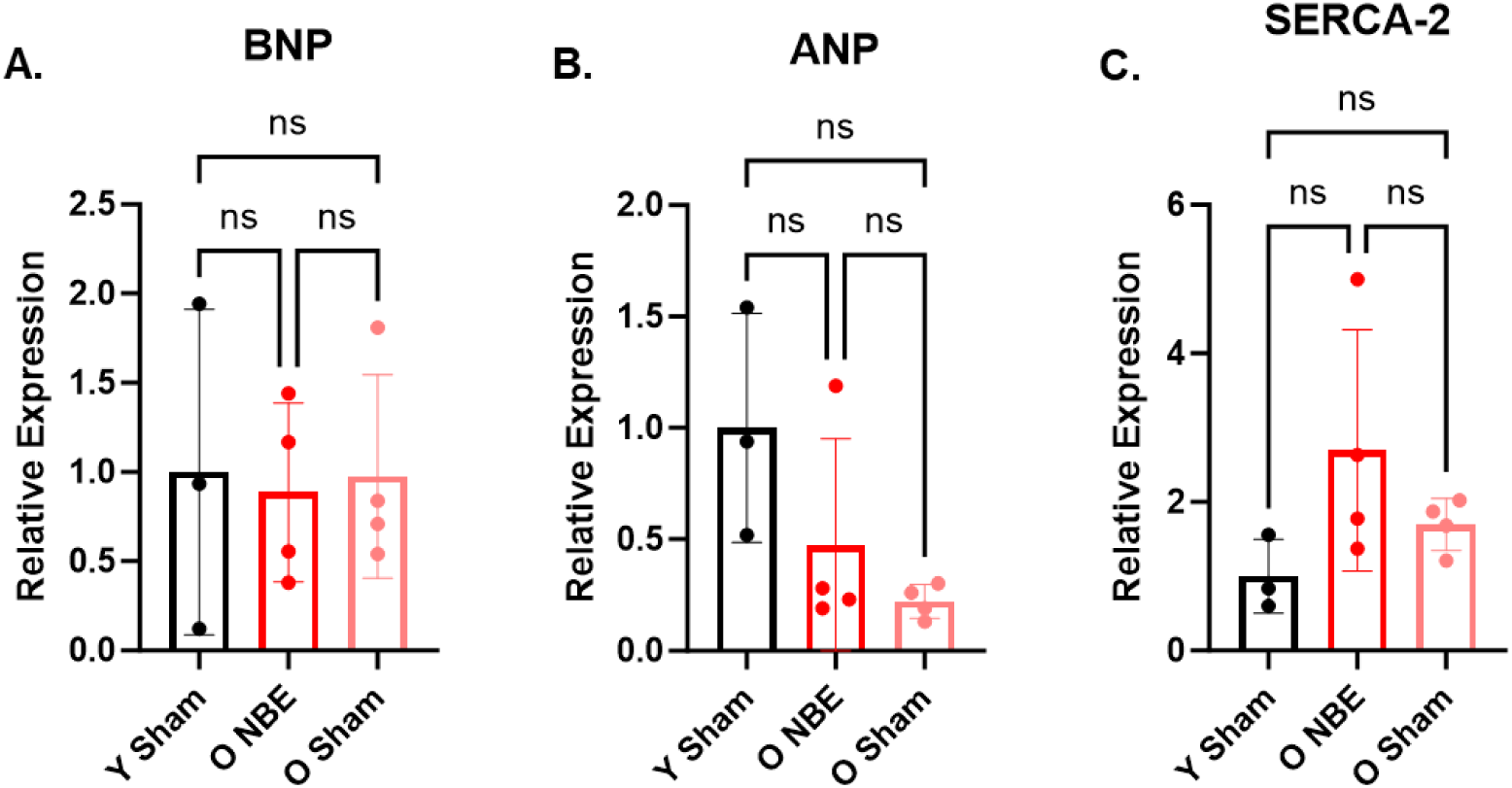
mRNA expression levels of brain natriuretic peptide, atrial natriuretic peptide, and SERCA-2 are highly variable. **A.** Relative transcript levels for BNP, ANP (**B**), and SERCA-2 (**C**). For each graph, n = 3 for Y sham and n = 4 for ONBE and O sham (all individual animals). Ns = non-significant by One-Way ANOVA, Tukey posthoc.

**Supplementary Figure 5.**
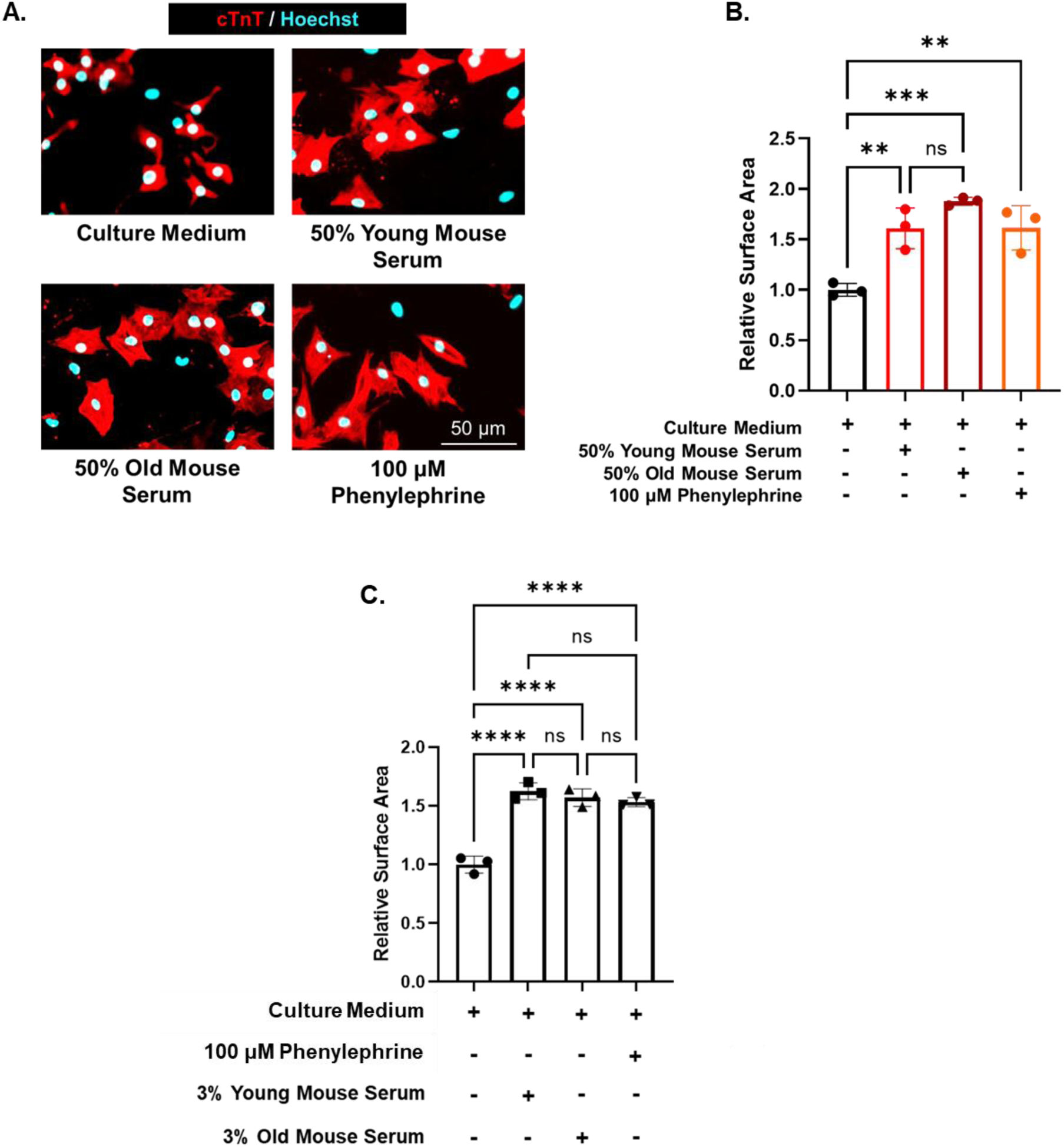
Young and aged mouse sera induce hypertrophy in neonatal rat cardiomyocytes. P2 rat cardiomyocytes (CMs) were freshly isolated in Dulbecco’s Modified Eagle Medium (DMEM) supplemented with 10% FBS and 20 μM cytosine arabinoside (AraC) for approximately 24 hours. Next, CMs were treated with the sera indicated above for an additional 24 or 48 hours. CMs were then fixed with 4% PFA and immunostained with anti-cTnT antibodies. Both young and aged sera induced CM hypertrophy to a similar extent. **A.** Representative images of P2 rat CMs after treatment with accompanying quantification (**B**). **C.** Quantification of P2 rat CMs after treatment with 3% young or 3% aged mouse sera. Relative surface areas were increased to amounts that were comparable to those of CMs treated with 50% sera. Data is shown as technical triplicates per cohort. ** P < 0.01, *** P < 0.005, **** P < 0.001 by One-Way ANOVA, Tukey posthoc. Scale bar = 50 μm. Data shown as mean +/-Stdev.

## Notes

### Competing Interest Statement

The authors have declared no competing interest.

